# Chk1 Phosphorylates Cdh1 to Promote SCF^βTRCP^-Dependent Degradation of Cdh1 During S-Phase

**DOI:** 10.1101/535799

**Authors:** Debjani Pal, Adrian E. Torres, Abbey L. Messina, Andrew Dickson, Kuntal De, Belinda Willard, Monica Venere, Matthew K. Summers

## Abstract

The interplay of the Anaphase-Promoting Complex/Cyclosome (APC/C) and Skp1-Cul1-F-box (SCF) E3 ubiquitin ligases is necessary for controlling cell cycle transitions and checkpoint responses, which are critical for maintaining genomic stability. Yet, the mechanisms underlying the coordinated activity of these enzymes are not completely understood. Recently, Cyclin A- and Plk1-mediated phosphorylation of Cdh1 was demonstrated to trigger its ubiquitination by SCF^βTRCP^ at the G1/S transition. However, Cyclin A-Cdk and Plk1 activities peak in G2 so it is unclear why Cdh1 is targeted at G1/S but not in G2. Here, we show that phosphorylation of Cdh1 by Chk1 contributes to its recognition by SCF^βTRCP^, promotes efficient S-phase entry, and is important for cellular proliferation. Conversely, Chk1 activity in G2 inhibits Cdh1 accumulation. Overall, these data suggest a model whereby the rise and fall of Chk1 activity is a key factor in the feedback loop between APC/C^Cdh1^ and the replication machinery that enhances the G1/S and S/G2 transitions, respectively.

## INTRODUCTION

Proper progression of the cell cycle is driven by the timely degradation of cell cycle regulators mediated by the ubiquitin proteasome system (UPS), which is necessary to maintain the systematic and coordinated duplication and subsequent segregation of the genome that is required to maintain its integrity [1-6]. Cells also possess a number of cell cycle checkpoints that work in conjunction with the UPS to further ensure orderly cell cycle progression and genome stability. Any defects in these processes may lead to irreversible damage including genetic alteration, developmental defects, and cancer [7].

There are over 600 different ubiquitin ligase in the human genome. Among them, the Skp1-Cullin-F-Box (SCF) complexes and the Anaphase Promoting Complex/Cyclosome (APC/C) are the best characterized for their roles in cell cycle control [8]. For SCF complexes, the F-box protein determines substrate specificity and substrate recognition by these proteins is often dependent on post-translational modification of the substrate. For the majority of the well-studied F-boxes (e.g., Skp2, Fbw7, βTRCP) it is substrate phosphorylation that allows efficient recognition by the F-box protein [2, 9]. Similar to the SCF, the APC/C activity and substrate recognition depends on either one of two WD40 repeat proteins, Cdc20 or Cdh1[10-15]. While APC/C^Cdc20^ participates almost exclusively in mitotic progression, the biological function of APC/C^Cdh1^ is much more complex and broad. The primary functions of APC/C^Cdh1^ include mitotic exit, G1 maintenance, quiescence, and differentiation [7, 16, 17]. Cdh1 plays key roles in the maintenance of chromosomal integrity and genomic stability [18-26]. In keeping with its diverse functions, Cdh1 has been established as a tumor suppressor as Cdh1-deficient mice exhibit genomic instability and develop epithelial tumors[21]. Indeed, many known Cdh1 substrates including Cyclin A, Plk1, Ect2, HEC1, Aurora kinases, and Skp2 are overexpressed in various cancers with high genomic instability and are associated with oncogenesis [16, 18, 21, 27]. Thus, dysregulation of APC/C^Cdh1^ could play a significant role in carcinogenesis.

The activity of APC/C^Cdh1^ is tightly regulated and must remain low for the G1 to S phase transition as some of its substrates participate in S-phase entry and the replication process. Multiple mechanisms work together to keep the activity of APC/C^Cdh1^ low from late G1 until late mitosis. For instance, degradation of the E2 enzymes UbcH10 and UBE2S, mediated by APC/C^Cdh1^ in G1, attenuates APC/C^Cdh1^ activity [28-30]. Accumulation of Emi1 inhibits APC/C^Cdh1^ by acting as a pseudosubstrate and preventing ubiquitin chain elongation [31-36]. As APC/C^Cdh1^ activity diminishes, the accumulation of substrates such as Cdc25A, Skp2 and Cyclin A promote phosphorylation of Cdh1 by Cyclin-Cdk complexes, which further weakens APC/C^Cdh1^ activity by disrupting the interaction of Cdh1 with APC/C [11, 28, 37]. Degradation of Cdh1 in late G1 is mediated by APC/C^Cdh1^ and SCF^Cyclin F^ [38, 39]. In addition, sequential phosphorylation of Cdh1 by both Cyclin A-Cdk2 and Plk1 leads to its degradation via the SCF^βTRCP^ E3 ubiquitin ligase [40]. Together these mechanisms cooperate to maintain low APC/C^Cdh1^ activity during S and G2 to ensure efficient cell cycle oscillation.

One question regarding the degradation of Cdh1 in late G1/S that still remains is why Cdh1 is targeted by SCF^βTRCP^ at this stage of the cell cycle when the activities of cyclin A-Cdk, and particularly PLK1, are at their lowest, but is not targeted in G2 when the activities of these kinases are maximal. We therefore speculated that an additional, S-phase active kinase would be involved. The Chk1 kinase was recently implicated in the regulation of Cdh1 following replication stress and several lines of evidence indicated Chk1 as a candidate Cdh1 kinase [41]. Chk1 is a downstream effector of the ATR kinase and is required for the cellular response to replication stress including promoting the SCF^βTRCP^-mediated degradation of Cdc25 [42, 43]. Chk1 is also central to the normal control of the replication program during S-phase and early activation of Chk1 is associated with premature S-phase entry, similar to loss of APC/C^Cdh1^ activity [44, 45]. Finally, APC/C^Cdh1^ is a negative regulator of Chk1 activitation, suggesting that a feedback loop between these proteins may exist [20, 44, 46, 47]. Herein, we determined that Chk1 phosphorylates Cdh1 promoting its recognition by SCF^βTRCP^ for ubiquitination and subsequent proteasome-mediated degradation. We further defined that expressing a constitutively active Chk1 prevents Cdh1 accumulation in G2 phase. Together our data provide a model whereby Chk1 activity in S-phase cooperates with Cyclin A-Cdk2 and low Plk1 activity to facilitate recognition of Cdh1 by SCF^βTRCP^ whereas loss of Chk1 activity in G2 permits Cdh1 accumulation despite increasing Plk1 and Cdk activity.

## RESULTS AND DISCUSSION

### Chk1 Modulates Cdh1 Stability

Recent findings by the Bartek group suggest that activated Chk1 signaling promotes degradation of Cdh1 [41]. However, whether Cdh1 might be acted upon by Chk1 to elicit these effects was not known. To address that possibility, we first examined the impact of Chk1 inhibition on Cdh1 stability by disrupting Chk1 with the Chk1 inhibitor, Chir124 (Figure 1A), or Chk1 siRNA (Figure 1B). Both asynchronously dividing HeLa and 293T cells showed an increase in the abundance of endogenous Cdh1 upon inhibition of Chk1. Consistent with previous work [41], depletion of Chk1 in asynchronous 293T cells significantly increased endogenous Cdh1 levels (Figure. 1B). Furthermore, as Chk1 inhibition will allow cells to progress to mitosis, where Cdh1 stability is increased, we examined Cdh1 levels in non-mitotic cells after Chk1 inhibition and also observed an increase in Cdh1 (Figure 1C). As an additional strategy to evaluate the relationship between Chk1 and Cdh1 stability, we exposed cells to hydroxyurea-induced replication stress as replication stress is known to reduce the half-life of Cdh1 [41, 48]. In agreement with a role for Chk1 in endogenous Cdh1 stability, we further found that inhibition of endogenous Chk1 leads to a significant increase in the steady state levels of both exogenously expressed Cdh1 (Figure 1D) and endogenous Cdh1 (Figure EV1) even in the presence of hydroxyurea-induced replication stress. Taken together, these data confirm the involvement of Chk1 in the regulation of Cdh1 protein abundance.

**Figure 1:**
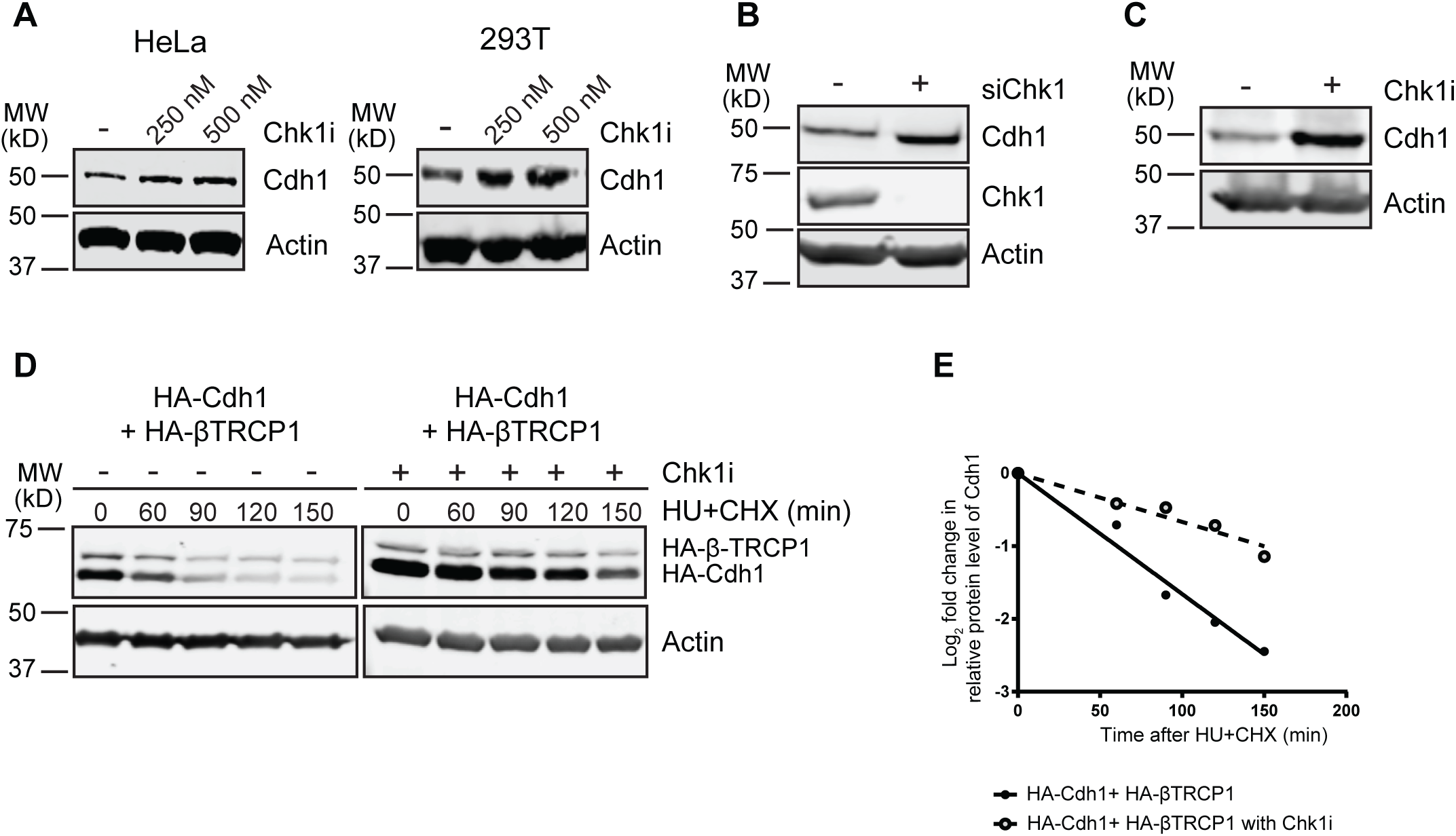
Chk1 Modulates Cdh1 Stability. **(A)** Inhibition of Chk1 increases the stability of endogenous Cdh1. Immunoblot analysis of whole-cell lysates derived from both HeLa and 293T cells treated with increasing doses of the Chk1 inhibitor, Chir124, for 5 hr before harvesting. **(B)** Depletion of endogenous Chk1 leads to increased levels of endogenous Cdh1. Immunoblot analysis of whole-cell lysates derived from 293T cells transfected with the indicated siRNA. **(C)** Mitotic population does not contribute to the increased endogenous Cdh1 level after Chk1 inhibitor treatment. Immunoblot analysis of whole-cell lysates derived from HeLa cells after shake off the mitotic population post Chk1 inhibitor treatment for 5 hr. **(D and E)** Inhibition of Chk1 increases the half-life of transfected Cdh1 in 293T cells. **(D)** Immunoblot analysis of whole cell lysates derived from 293T cells, transfected with HA-Cdh1 and HA-βTRCP1 constructs. Cells were treated with both hydroxyurea (HU) and Chk1 inhibitor Chir124 (500 nM) for another 4 hr before addition of 50 µg/ml cycloheximide (CHX). At the indicated time points, whole-cell lysates were prepared for immunoblot analysis. **(E)** Quantification of the band intensities in **(D).** Cdh1 band intensities were normalized to actin and then further normalized to t=0 controls.

### SCF^βTRCP1^ Negatively Regulates Cdh1 at the G1-S Boundary in a Chk1 Dependent Manner

Fukushima et al previously reported that Cdh1 could be targeted by SCF^βTRCP^ in a CyclinA-Cdk2 and Plk1-dependent manner to promote the G1/S transition [40]. However, as both kinases are not highly active during the G1/S transition, when Cdh1 stability is low, but are highly active in G2, when Cdh1 stability is increased, we speculated that an additional S-phase-active kinase might cooperate with Cyclin A-Cdk2 and Plk1 to destabilize Cdh1. Chk1 plays a key role in the S-phase checkpoint and the replication process, suggesting that Chk1 as a candidate kinase that may contribute to targeting of Cdh1 by SCF^βTRCP^[49-58]. Given that replication stress promotes APC/C^Cdh1^ inactivation in a Chk1 dependent manner, we examined the role of Chk1 on the association of Cdh1 with βTRCP1. Pretreatment of HeLa G1/S extracts with the Chk1 inhibitor reduced the binding between GST-Cdh1 and HA-βTRCP1 (Figure 2A). Using an *in-vitro* kinase assay, we showed that phosphorylation of GST-Cdh1 with purified Chk1 served as a binding signal for βTRCP1 (Figure 2B). Together these data support that Cdh1 is targeted for βTRCP1-induced degradation in a Chk1-dependent fashion. To extend these *in-vitro* findings we determined the impact of Chk1 activity on the interaction of HA-Cdh1 and Flag-βTRCP1 in 293T cells. Co-expression of Myc-Chk1 increased the interaction between Cdh1 and βTRCP1 (Figure 2C) whereas inhibition of endogenous Chk1 had a negative impact on the association (Figure 2D). Thus, our biochemical data implicates a critical role for Chk1 mediated phosphorylation of Cdh1 in its recognition by βTRCP1 and subsequent degradation. Notably, these results are in keeping with previous research demonstrating that prior phosphorylation of Cdh1 is critical for it’s recognition by βTRCP1 via the DDGDVS sequence, which closely resembles the canonical DpSGx(2-4)pS degron sequence (where pS designates phosphorylated Ser residues) [59-63].

**Figure 2:**
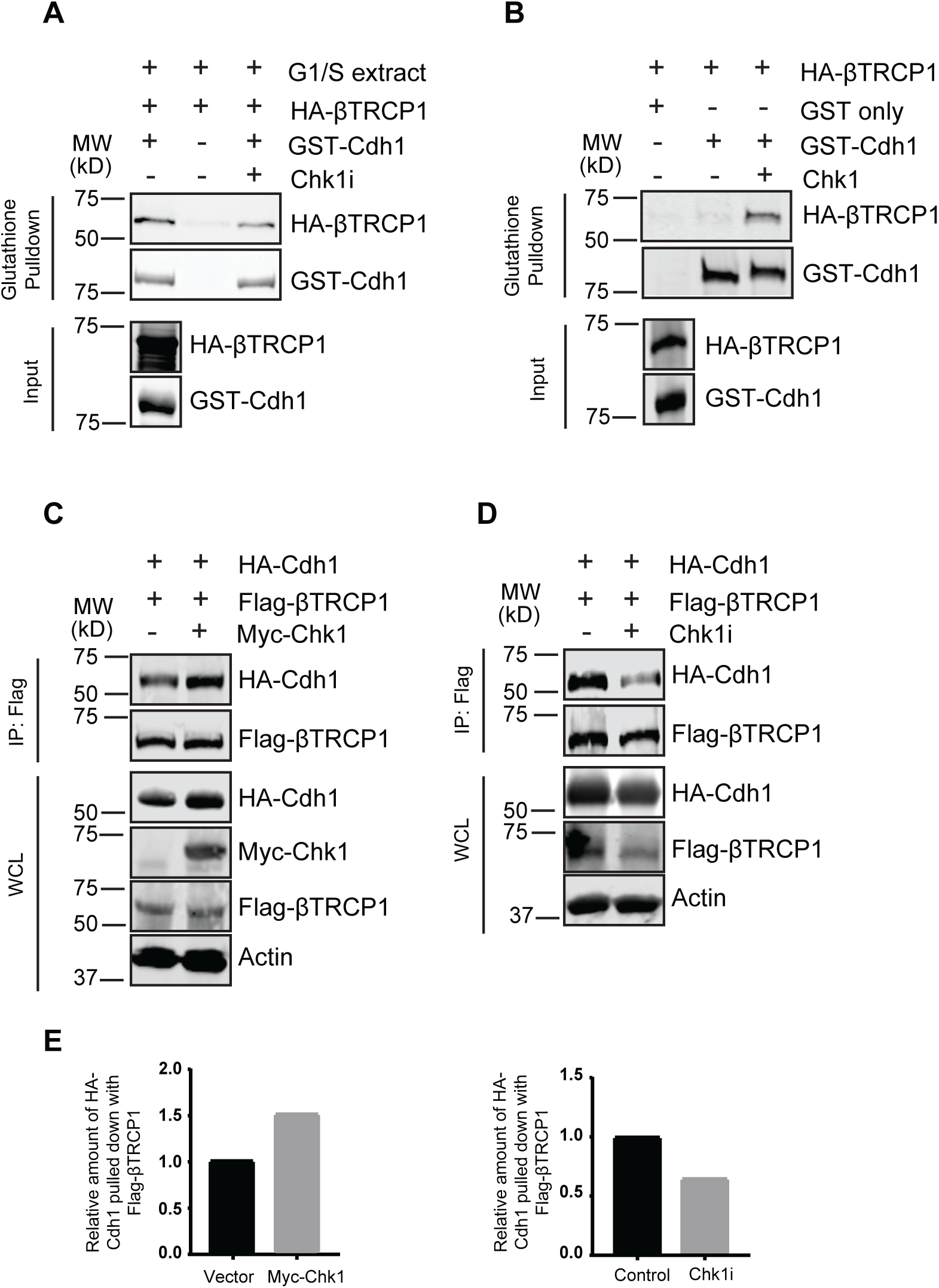
SCF^βTRCP1^ Negatively Regulates Cdh1 at the G1-S Boundary in a Chk1 Dependent Manner. **(A and B)** Chk1 impacts on the interaction between βTRCP1 and Cdh1 both in G1-S extracts and *in-vitro*. **(A)** HeLa cells were treated with HU (2 mM) for 20 hr to arrest them in G1-S boundary and extracts were prepared. The G1-S extracts were treated with Chk1 inhibitor where indicated before incubating with GST-Cdh1. After phosphorylation, GST-Cdh1 was first bound by glutathione beads and then mixed with *in-vitro* translated HA-βTRCP1 for an hour. Immunoblot analysis was carried out to detect the bound HA-βTRCP1 protein. **(B)** Immunoblots showing that the *in-vitro* phosphorylation of GST-Cdh1 by Chk1 promotes its binding with HA-βTRCP1. **(C and D)** Chk1 activity is crucial for the interaction of Cdh1 with βTRCP1 *in-vivo*. **(C)** Immunoblot analysis of immunoprecipitates and whole-cell lysates derived from 293T cells transfected with Flag-βTRCP1, HA-Cdh1 and Myc-Chk1 (where indicated). 30 hr post-transfection, cells were treated with proteasome inhibitor MG132 (10 µM) for 5 hr and then harvested to do immunoprecipitation to detect the interaction between HA-Cdh1 and Flag-βTRCP1 proteins. **(D)** Immunoblot analysis of immunoprecipitates and whole-cell lysates derived from 293T cells transfected with both Flag-βTRCP1 and HA-Cdh1. 30 hr post-transfection, Cells were treated with Chk1 inhibitor (where indicated) and proteasome inhibitor MG132 (10 µM) for 5 hr before harvesting to do IPs. **(E)** Quantification of the band intensities in **(C).** Immunoprecipiated HA**-**Cdh1 band intensities were normalized to their respected Flag-βTRCP1 IP bands and then further normalized to no Myc-Chk1 control set. **(F)** Quantification of the band intensities in **(D).** Immunoprecipiated HA**-**Cdh1 band intensities were normalized to their respected Flag-βTRCP1 bands and then further normalized to no Chk1 inhibitor treated control set.

### Phosphorylation of Cdh1 by Chk1 Creates a Phosphodegron Recognized by SCF^βTRCP1^

Given that inhibition of Chk1 promotes Cdh1 stability in response to DNA damage [41] and that Chk1 positively impacts the binding between Cdh1 and βTRCP1 both *in-vitro* and *in-vivo*, we explored the Chk1 phosphorylation event on Cdh1 more closely by an *in-vitro* kinase assay and mass-spectrometry. We first confirmed that Cdh1 could be directly phosphorylated by Chk1 (Figure 3A). The phosphorylated GST-Cdh1 was not detected in the presence of Chk1i or after phosphatase treatment (Figure 3A). An overall minimal consensus motif for Chk1 substrate phosphorylation is R/K-x-x-S/T. Intriguingly, S138, which was previously implicated in recognition of Cdh1 by βTRCP1, matches this consensus. However, although peptides containing phosphorylation of S138 and S137 could be identified by mass spectrometry analysis, these residues are not an efficient site for Chk1 mediated phosphorylation *in vitro* (not shown). In contrast, peptides containing phosphorylation at S133, S131/S133, and S172 were abundant with ratios of modified to unmodified peptides of ∼1, 20 and 34, respectively (Figure 3B and Appendix figure S1). Indeed, S172 conforms to the Chk1 consensus while S131/133 does not. Notably, the Phosphosite database (www.phosphosite.org) contains multiple examples of Chk1 substrates with non-conventional phosphorylation sites. We first decided to examine the possible involvement of those sites to impact βTRCP1 binding by introducing mutation of these serines to alanine. Given the previous implication of S138 we included S137 and S138 mutants in our studies. The interaction between Cdh1 and βTRCP1 was reduced by serine-to-alanine replacements of the Chk1 phosphorylation sites that we identified in Cdh1 (Figures 3C and EV2A). These data indicate that Cdh1 phosphorylation is a key step to induce βTRCP1 binding.

**Figure 3:**
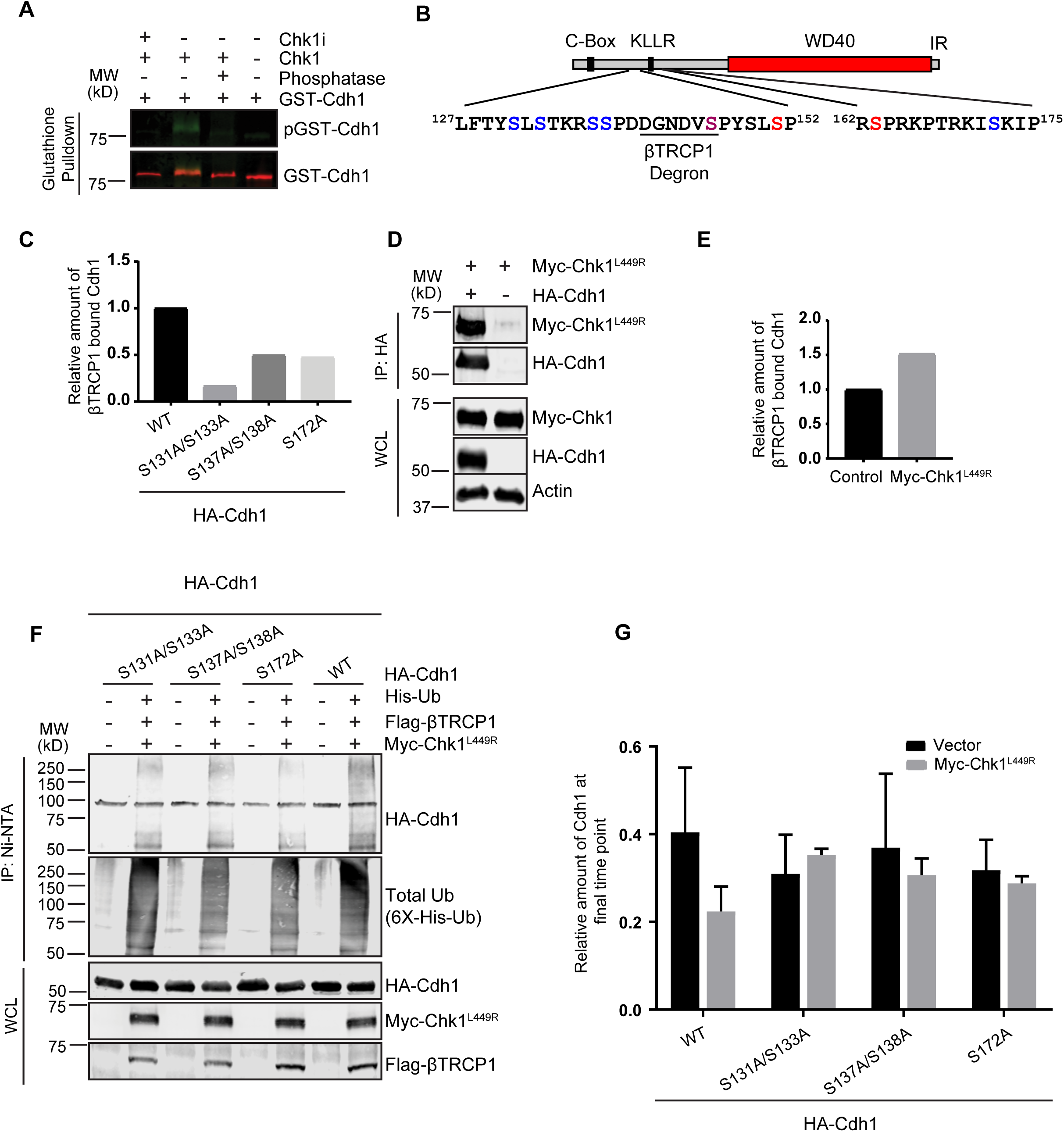
Phosphorylation of Cdh1 by Chk1 Creates a Phosphodegron Recognized by SCF^βTRCP1^. **(A)** Chk1 phosphorylates Cdh1 *in-vitro*. GST-Cdh1 was phosphorylated with Chk1 through *In-vitro* kinase assays. Immunoblot represents immunoprecipitated GST-Cdh1 on glutathione beads following in-vitro kinase assays. PIMAGO western blot kit was used to detect phospho-GST-Cdh1. Phosphorylation was further confirmed by including Chk1 inhibitor where indicated during the kinase assays or by treating phosphorylated GST-Cdh1 with phosphatase. **(B)** Schematic diagram of Cdh1. Chk1-mediated phosphorylation sites identified from mass-spectrometry analysis are indicated in blue. Cdh1 phosphorylation sites mediated by Cyclin A-Cdk2 and Plk1 are in red and magenta, respectively. The SCF-βTRCP1 phosphodegron is indicated. **(C).** Chk1 mediated phosphorylation of Cdh1 creates a binding site for βTRCP1. 293T cells were transfected with the indicated HA-Cdh1 constructs together with Flag-βTRCP1. Cells were treated with proteasome inhibitor MG132 (10 µM) for 5 hr and the interaction between HA-Cdh1 and Flag-βTRCP1 was analyzed. Immunoprecipiated HA**-**Cdh1 WT and the mutant band intensities were first normalized to their relative amounts in lysates. The relative amount of the indicated Cdh1 proteins bound to Flag-βTRCP1 was normalized to wild type HA-Cdh1. See also Figure **EV2A**. **(D)** Cdh1 interacts with Chk1 in vivo. Both HA-Cdh1 and constitutively active Myc-Chk1^L449R^ were co-expressed in 293T cells. 30 hr post-transfection, cells were treated with proteasome inhibitor MG132 (10 µM) for 5 hr. The interaction between HA-Cdh1 and Myc-Chk1^L449R^ proteins was monitored by immunoprecipitation. **(E)** Constitutively active Chk1 promotes Cdh1 and SCF**^βTRCP1^** interaction *in-vivo*. Immunoblot analysis of immunoprecipitates and whole-cell lysates derived from 293T cells transfected with Flag-βTRCP1, HA-Cdh1 and Myc-Chk1^L449R^ (where indicated). 30 hr post-transfection, cells were treated with proteasome inhibitor MG132 (10 µM) for 5 hr. Immunoprecipiated HA**-**Cdh1 band intensities **(EV2B)** were normalized to their respected Flag-βTRCP1 IP bands and then further normalized to control. **(F)** *n-vivo* ubiquitination assays shows that SCF**^βTRCP1^** promotes Cdh1 ubiquitination in a Chk1 dependent manner. **(G)** Mutations of Chk1 mediated phosphorylation sites in Cdh1 increases the stability of Cdh1. 293T cells were transfected with the indicated HA-Cdh1 constructs together with Flag-βTRCP1 and Myc-Chk1^L449R^ (where indicated). Cells were treated with 50 µg/ml cycloheximide (CHX). At the indicated time points, whole-cell lysates were prepared for immunoblot analysis. The intensities of Cdh1 bands were normalized to actin, then normalized to the t=0 time point. The plots represent the relative fraction of indicated Cdh1 protein at final time point (t=320 min) after adding CHX. Data are represented as mean +/- SD, n=3 biological replicates. See also **(EV2C)**.

Chk1 exists in a closed, inactive confirmation [64]. Upon replication stress or DNA damage, ATR-dependent phosphorylation of Ser-317 and Ser-345 activates Chk1 by antagonizing intramolecular interactions that stabilize the closed confirmation and making the Chk1 catalytic site available to interact with substrates [49, 55, 64]. Introduction of the L449R mutation disrupts the closed conformation, resulting in a constitutively active Chk1 [64]. We utilized this constitutively active Chk1^L449R^ to test the impact of Chk1 activity on Cdh1 degradation in the absence of additional replication stress. Cdh1 interacts with Chk1^L449R^ when expressed together in 293T cells (Figure 3D). In agreement with a role for Chk1 in Cdh1 and βTRCP1 interaction, we further found that co-expression of Chk1^L449R^ with Cdh1 and βTRCP1 led to increased HA-Cdh1 binding with Flag-βTRCP1 (Figures 3E and EV2B) *in vivo*. The Cdh1 degron DDGNDVpS is non-canonical. However, such sites often have additional phosphorylation sites surrounding them. To examine the possible involvement of those sites for βTRCP1-dependent ubiquitination of Cdh1, *in vivo* ubiquitination assay were carried out with each of the Chk1 phosphorylation site mutants of Cdh1. Ubiquitination was greatly reduced with the alanine mutants of Cdh1 compared to wild type even in the presence of active Chk1 (Figure 3F). In agreement these data, co-expression of βTRCP1 and Chk1^L449R^ significantly reduce the stability of wild type Cdh1 in the presence of cycloheximide whereas S131/133A, S137/138A and S172A mutant Cdh1 were not impacted by the presence of Chk1^L449R^ (Figures 3G and EV2C). Overall, these data established that Chk1 dependent phosphorylation of Cdh1 is important for its recognition by βTRCP1 and subsequent degradation. This phosphorylation could be a part of Chk1 mediated G1/S surveillance mechanism where downregulation of Cdh1 is required to establish the replication process. This data also supports our hypothesis that multiple kinase work together to regulate Cdh1 level for proper cell cycle transition.

### Maintaining Chk1 Activity Inhibits the Accumulation of Cdh1 Through S/G2 Phase

We established that Chk1 mediated phosphorylation promotes Cdh1 degradation by βTRCP1 along with CyclinA-Cdk2 and Plk1 in G1/S phase. In contrast to high levels of Cyclin A-Cdk2 and Plk1 activities, multiple mechanisms cooperate to attenuate Chk1 activity in G2 [46, 52, 65-69]. We therefore hypothesize that in G2 the absence of Chk1 activity facilitates Cdh1 stability and accumulation. Expression of constitutively active Chk1^L449R^ in synchronized 293T cells significantly reduced endogenous Cdh1 accumulation through S/G2 upon release from HU treatment (Figures 4A and 4B). The cells progressed through S/G2 phase normally as confirmed by flow cytometry analysis (Figure 4C). This data identifies a unique mechanism for allowing Cdh1 stability in G2 phase of the cell cycle.

**Figure 4:**
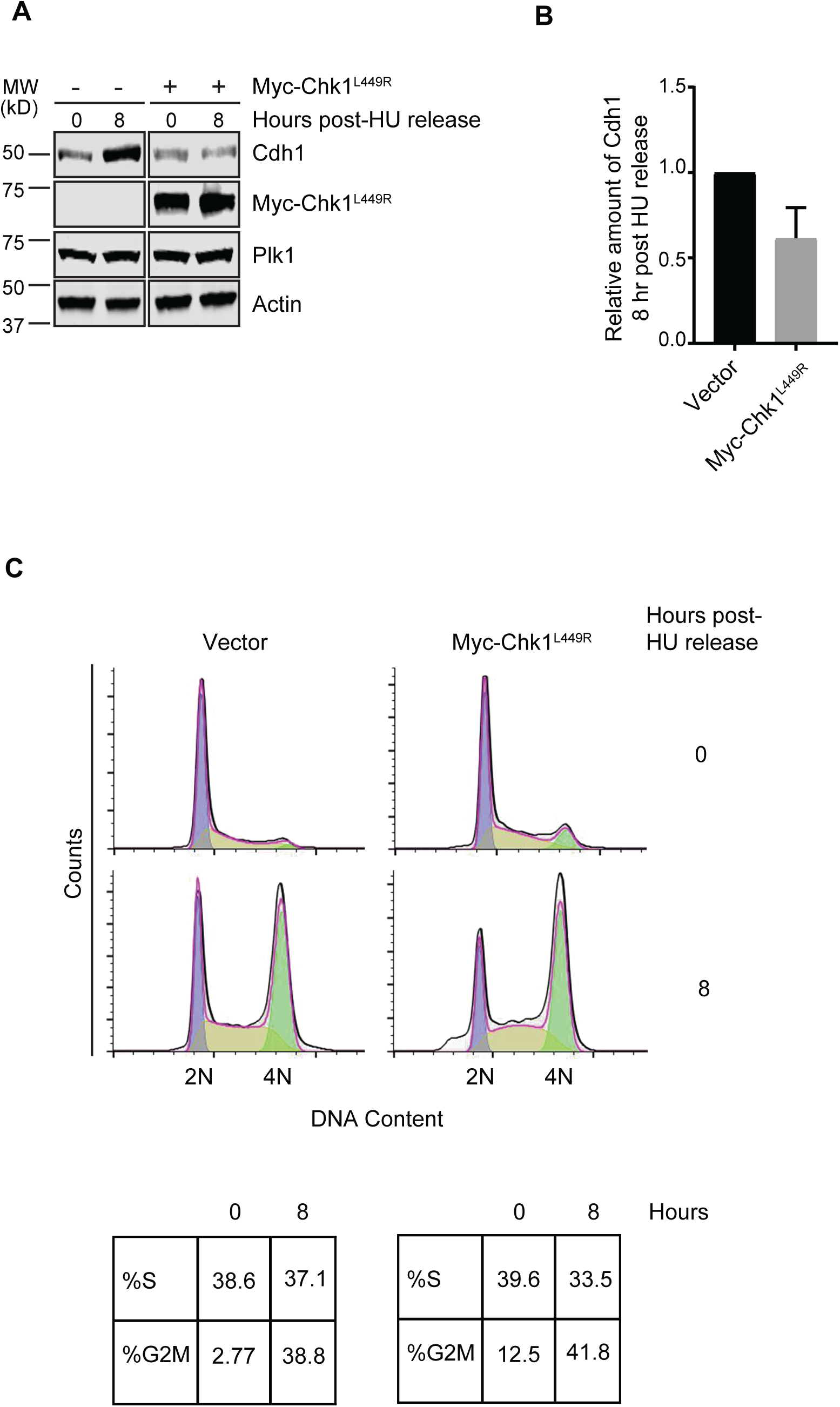
Presence of Constitutively Active Chk1, Inhibits the Accumulation of Cdh1 Through S/G2 Phase. **(A-C**) Immunoblot analysis of whole-cell lysates of 293T cells transfected with Myc-Chk1^L449R^. **(A)**. Cells were treated with HU (2 mM) for 20 hr and released into the fresh media. At the indicated time points, cells were harvested to prepare the whole-cell lysates for immunoblot analysis. The experiments were performed twice. **(B)** Quantification of band intensities as in **(A).** The intensities of Cdh1 were normalized to actin and then further normalized to vector control. Data are represented as mean +/- SD, n=2 biological replicates. **(C)** Cell cycle progression was monitored by flow cytometry. The experimental set up was as described as in 4A.

### Chk1-Mediated Phosphorylation of Cdh1 Destabilizes the Interaction between Cdh1 and the APC/C

We next considered the possibility that the Chk1 phosphorylation sites may regulate the binding of APC/C and Cdh1. The APC/C-Cdh1 interaction is known to be regulated by the phosphorylation status of Cdh1. Cyclin-Cdk mediated phosphorylation of Cdh1 has an adverse impact on APC/C^Cdh1^ assembly and prevents any unscheduled activation of APC-Cdh1 complex [11, 28, 37, 60, 70, 71]. The Chk1 phosphorylation sites on Cdh1 are in proximity to the APC/C binding interface of the Cdh1 C-box and KLLR motifs and could potentially influence its interaction with the holoenzyme[72]. To further explore the possibility the Chk1-mediated phosphorylation may peturb APC/C^Cdh1^ complex formation, *in-vitro* translated Cdh1 proteins were incubated with G1/S HeLa cell extracts treated with or without Chk1i and the ability of Cdh1 to interact with the APC/C was examined by co-immunoprecipitated of Cdh1 with the APC/C. Consistent with previous reports, wild type Cdh1 showed minimal binding to APC/C[24, 28, 73]. However, APC/C-Cdh1 binding was strongly increased when Chk1 was inhibited. Notably, Cdh1 phosphosite mutants showed higher basal APC/C binding that was not increased by Chk1i (Figure EV3). These data open up an entirely new role for Chk1 kinase in regulating Cdh1 activity by not only promoting its downstream ubiquitination but also destabilizing its association with the APC/C complex itself to maintain low Cdh1 activity during replication.

### Chk1 Mediated Phosphorylation of Cdh1 Ensures Proper Cell-Cycle Progression

Given that mutation of the Chk1 phosphorylation sites of Cdh1 extended the half-life of Cdh1 and considering the importance of Cdh1 inactivation for S-phase entry, we next sought to determine the physiological significance of Chk1 mediated Cdh1 phosphorylation on cell growth. HeLa cells were transfected with both wild type and Cdh1 phosphosite mutants along with H2B-GFP and proliferation was monitored in real time. All the cells expressing non-degradable Cdh1 mutants showed a dramatic reduction in proliferation in comparison to control and wild type-expressing cells (Figure 5A). In addition, we observed an increased number of cells with enlarged nuclei in the populations expressing the phosphosite mutants in comparison to wild type and control cells (Figure 5B and Appendix Figure S2A). This observation is consistent with the re-replication of cellular DNA caused by aberrantly increased APC/C^Cdh1^ activity [24, 28, 74-77]. To specifically address the role of Chk1-mediated phosphorylation of Cdh1 on S-phase entry, we monitored the progression of cells through G1 and into S-phase following release from a mitotic arrest. Cells were pulse labeled with EdU to identify replicating cells 8, 10, and 12 hours after release from the mitotic block. Analysis of EdU-positive cells revelaed that cells expressing Cdh1 phosphorylation site mutants exhibit significantly delayed S-phase entry compared to vector control or wild type Cdh1-expressing cells (Figure 5C and Appendix Figure S2B).

**Figure 5:**
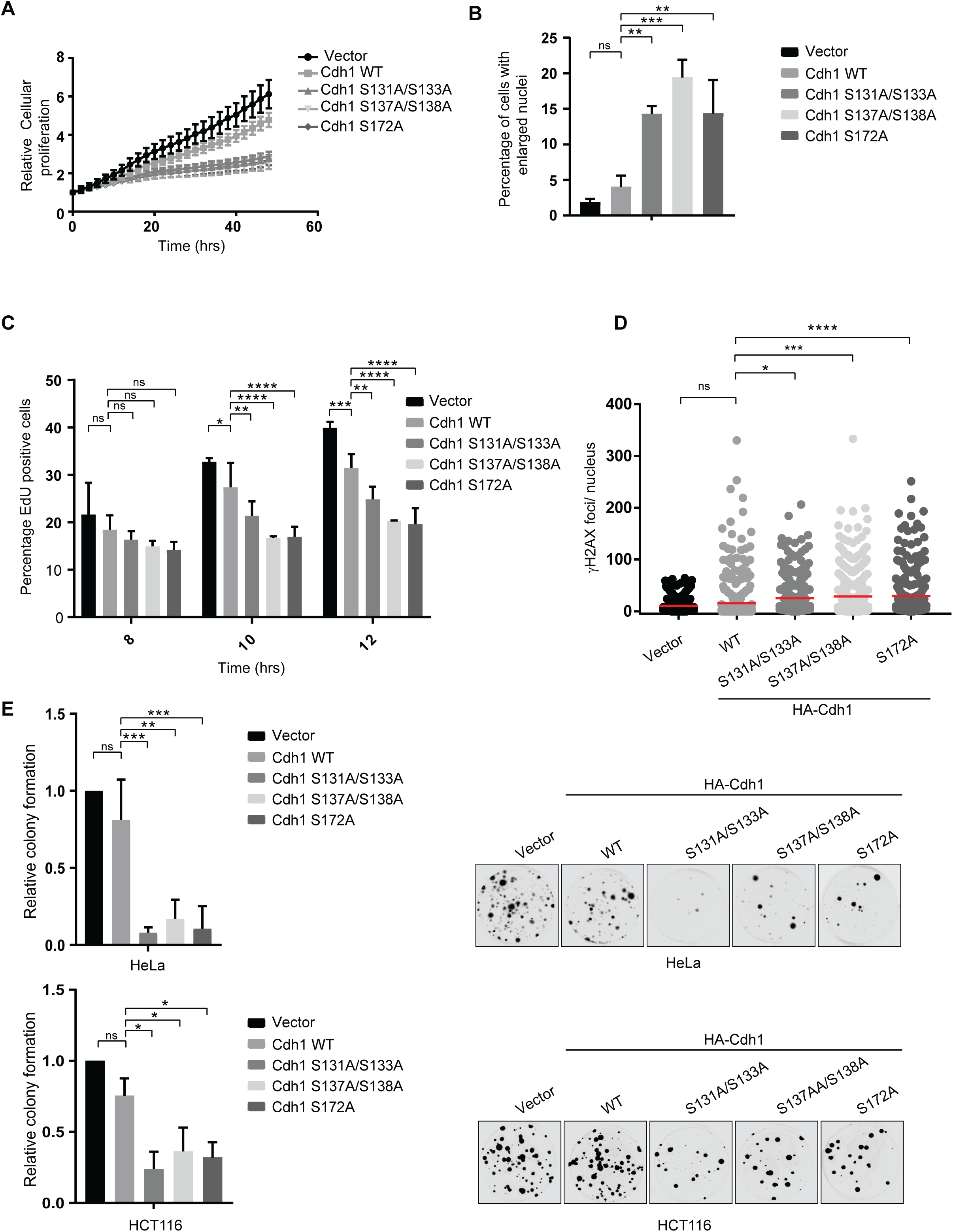
Chk1 Mediated Phosphorylation of Cdh1 Ensures the Proper Cell-Cycle Progression. **(A-C**) Mutation of the Chk1 mediated Cdh1 phosphorylation delays S-phase progression. HeLa cells were transfected with indicated Cdh1 constructs and histone H2B-GFP. After 24 hours, proliferation **(A)** and nuclear size **(B)** of the GFP-positive nuclei were analyzed. **(B)** The figure represents the percent of GFP positive cells with enlarged nuclei. Data **(A and B)** are represented as mean +/- SD, n=3 biological replicates **p<0.01, ***p<0.001 were calculated with 1-way Anova with Sidak’s post-test. **(C)** Hela cells were synchronized in mitosis with a thymidine-nocodazole block following transfected with the indicated Cdh1 constructs. After release from nocodazole, cells were pulsed with Edu for 15 min, at the indicated time points fixed and analyzed for Edu incorporation. The plot represents the percent of GFP-positive, Edu-positive nuclei. Data are represented as mean +/- SD, n=3 biological replicates and *p< 0.05, **p<0.01, ***p<0.001, ****p<0.0001 were calculated with 2-way Anova with Dunnet’s post-test. **(D)** Mutation of the Chk1 phosphorylation sites in Cdh1 induces DNA damage. The number of γH2AX foci in cells. U2OS cells as in (A) were analyzed for γH2AX foci. The graph shows the number of γH2AX foci per GFP-positive nucleus, n>275; *p< 0.05, ***p<0.001, ****p<0.0001 were calculated with 1-way Anova with Dunnet’s post-test. **(E)** Effects of non-degradable Cdh1 on colony-forming capacity of HeLa and HCT116 cells. Cells were co-transfected with the indicated Cdh1 constructs. The graph represents the relative fraction of number of colonies transfected with each indicated Cdh1 constructs and normalized to vector control. n=3 biological replicates (with three technical replicates per biological replicate), error bars represents standard deviation and *p< 0.05, **p<0.01, ***p<0.001 were calculated with 1-way Anova with Sidak’s post-test. Representative images of colony formation following each treatment are shown for both HeLa and HCT116 cells.

To investigate the nature of the cell cycle delay during G1 phase observed after overexpression of non-degradable Cdh1, cells were examined by immunofluorescence analysis for γH2AX foci and 53BP1 foci, markers of DNA damage. Cells transfected with Cdh1 mutants showed dramatically increased counts for these foci indicating a higher level of DNA damage (Figures 5D, Appendix Figure S2C, and Appendix Figure S3). In addition, clonogenic survival assay showed that stabilization of Cdh1 significantly lowered cellular proliferation for both HeLa and HCT116 cells (Figure 5E). Our data demonstrate that Chk1 phosphorylation plays a significant role, along with previously demonstrated functions of Plk1 and CyclinA-Cdk2, to promote SCF^βTRCP1^ mediated downregulation of Cdh1 for timely S phase entry and to enhance the maintenance of genomic stability.

Overall, our study has identified Cdh1 as a previously unknown substrate of the Chk1 kinase. This finding provides new insight into the regulation of Cdh1 by Chk1 and SCF^βTRCP^. First, we have identified Chk1 phosphorylation of Cdh1 in the regions flanking the SCF^βTRCP^-recognized phosphodegron as an important facet in the recognition of Cdh1 by the ubiquitin ligase. Consistent with this idea, exposure to UV radiation, a potent inducer of Chk1 activity, triggers Cdh1 degradation [48]. Although the role of Chk1 and SCF^βTRCP^ were not formally examined, the degron-containing region of Cdh1 identified in these studies contains the SCF^βTRCP^ phosphodegron and the Chk1 phosphorylation sites, supporting a potential role for the kinase in these events. Previous work had identified Plk1 as the kinase responsible for phosphorylation of the core degron at S146 and demonstrated that phosphorylation of the region flanking the core degron as important for the interaction between SCF^βTRCP^ and Cdh1, specifically S138 [40]. Given that Plk1 activity increases during G2, the involvement of an S-phase-active kinase (e.g., Chk1) in the recognition of Cdh1 by SCF^βTRCP^ provides a mechanism by which Cdh1 protein is able to accumulate during this phase [78]. Indeed our data suggest that down-regulation of Chk1 activity promotes Cdh1 accumulation. Rising Plk1 activity in G2 weakens Chk1 activation by triggering the destruction of Claspin, while active Chk1 is targeted by the SCF^Fbx6^ and CRL4^Cdt2^ ligases[65-69, 79, 80]. As Chk1 activity diminishes, Cdh1 stability increases allowing it to target Rad17 further weakening Chk1 activity and contributing to a feedforward loop to promote cell cycle progression beyond S-phase[46]. In contrast, overexpression of Claspin leads to Chk1 activation and early S-phase entry as well, suggesting that feedback toward Cdh1 via the replication checkpoint machinery sharpens the G1/S transition (Figure EV4)[44]. In keeping with this idea the deubiquitinating enzyme USP37, which we have previously shown to facilitate S-phase entry, stabilizes Chk1 (manuscript in preparation) and cells depleted of USP37 demonstrate attenuated Cdh1 degradation in late G1 and delayed Chk1 phosphorylation (Figure EV5) as well as delayed initiation of replication[81]. However, further studies are required to determine whether Chk1-mediated inhibition of APC/C^Cdh1^ plays a role in the ability of UPS37 to promote S-phase entry.

From a mechanistic standpoint, the region surrounding S138 resembles the general Chk1 consensus site and Chk1 has recently been demonstrated to promote Cdh1 degradation upon replication stress. However, although mutation of S137 and 138 rendered Cdh1 resistant to Chk1-enhanced degradation, we did not obtain evidence for robust direct phosphorylation of these residues by Chk1. We interpret this result to indicate that Chk1 may indirectly regulate the phosphorylation of these residues as has been demonstrated for Cdc25A[82]. However, we cannot exclude the possibility that S137, S138 of our recombinant Cdh1 is simply not available to the kinase.

In contrast to S137, S138, Chk1 efficiently catalyzed the direct phosphorylation of S131, S133, and S172 and mutation of these sites abolished the ability of Chk1 to promote Cdh1 degradation. These sites are poorly characterized, with a single reported proteomic identification for S131 and S133 and no previous reports of S172 phosphorylation. Intriguingly, increased phosphorylation of these residues is observed in glioblastoma cancer stem cells (De et al, manuscript submitted), which we have previously shown to have attenuated Cdh1 activity. Phosphorylation at S131/133 is well positioned to contribute to the recognition of CDH1 by βTRCP as supported by the ability of Chk1 to promote binding between Cdh1 and βTRCP in vitro. The contribution of S172 phosphorylation is less clear as the distance from the phosphodegron would suggest that this site is less likely to contribute to βTRCP-binding. From a structural standpoint, S172 appears to reside at the C-terminal end of a linker between the APC/C interacting KLLR motif and the WD40 domain and may pack against the WD40 propeller[72]. Thus, phosphorylation of this residue could impact the stability of the Cdh1 structure and/or the interaction of Cdh1 with APC/C, which may promote the availability of Cdh1 to additional kinases as well as βTRCP. Indeed our data indicate that phosphorylation of Cdh1 by Chk1 regulates the interaction with APC/C. This finding is in line with previous work in fission yeast where the replication stress checkpoint effector, Cds1, negatively regulates APC/C activity by phosphorylating the Cdh1 homolog Ste9 and preventing its interaction with the holoenzyme [83]. Moreover, while phosphorylation of these sites is important for regulation of APC/C^Cdh1^ activity and cellular well-being, mutation of S133 and S172 have been identified in tumors (www.cbioportal.org) Interestingly, both residues are subject to S→F mutations, suggesting that despite an inability to regulate Cdh1 by phosphorylation the presence of the bulky Phe side-chain may negatively regulate the APC/C-Cdh1 interaction and circumvent the deleterious effects of failing to down-regulate Cdh1. The ability of Chk1 to antagonize APC/C-Cdh1 interaction provides further insight into the checkpoint recovery switch proposed above. As Chk1 activity diminishes, Cdh1 would not only become less well recognized by βTRCP, but would also acquire an enhanced ability to bind the APC/C to prevent further Chk1 activation and facilitate its accumulation.

Given that loss of APC/C^Cdh1^ activity is associated with genomic instability and tumorigenesis it is tempting to speculate that upregulation of Claspin, Chk1, Cyclin A, and Plk1 may contribute to genomic instability and cancer, in part, via antagonism of APC/C^Cdh1^. In contrast, while multiple mutations in *FZR1*, the gene encoding Cdh1, across several tumor types cluster in the degron region of the protein, the majority of these mutations (e.g., D140N, D144N, within the core degron) would seem likely to antagonize down-regulation of Cdh1 rather than promote it (www.cbioportal.org). Our observations of inefficient initiation of replication, increased DNA damage, and evidence of genomic instability in the form of DNA re-replication in cells that failed to down-regulate Cdh1 is in direct line with previous studies showing that down-regulation of Cdh1 at the G1/S transition and during S-phase, particularly in the presence of replication stress, is critical for cellular viability and genomic stability. Notably, the more conservative mutations in non-phosphorylated sites within this region may have a less severe impact on the degradation of Cdh1 than ablation of Cdh1 phosphorylation sites used in this and previous studies. Thus, we postulate that future examination of these tumor-derived mutations will reveal increased genomic instability, but at a level compatible with viability, and provide further evidence that altered regulation of APC/C^Cdh1^, either positive or negative, has pathophysiological consequences.

## Materials and Methods

### Mammalian Cell Culture, synchronization, and drug treatments

HeLa, 293T and HCT116 cells were obtained from ATCC and maintained in DMEM complete medium (Corning) supplemented with 10% fetal bovine serum (FBS; Seradigm). Hela cells were synchronized in S-phase by double thymidine block with 2 mM thymidine and transfection between the blocks followed by treatment with 50 ng/ml nocodazole to arrest cells in mitosis. The cells were then washed two times with fresh DMEM complete medium and replated into nocodazole-free fresh medium. For USP37 siRNA treatment, HeLa cells were transfected with RNAiMAX (Invitrogen) per the manufacturer’s instructions between double thymidine. 293T cells were arrested at G1/S phase in 2 mM hydroxyurea for 16 hr and washed with PBS and released into fresh DMEM complete medium. Plasmid transfections were done with TransIT-LT1 (Mirus Bio) per the manufacturer’s instructions. Where indicated, cells were treated with 500 nM CHIR-124 (Selleckchem), 200 nM BI2536 and 10 µM MG132 (Boston Biochem). Cycloheximide (CHX) assay was performed as described previously.

### Immunofluorescence, microscopy, and flow cytometry

were performed as previously described [84]. Detection of DNA synthesis in proliferating cells was determined based on the incorporation of 5-ethynyl-2′-deoxyuridine (EdU; Thermo Fisher Scientific) and its subsequent detection by a fluorescent azide through a Cu(I)-catalyzed [3 + 2] cycloaddition reaction (“click” chemistry) per the manufacturer’s instructions. In brief, HeLa cells were transfected with different Cdh1 constructs and histone H2B-GFP as a tracer and synchronized in S-phase with double treatment with 2 mM thymidine and then arrested at G2/M phase in 50 ng/ml Nocodazole for 16 hr treatment. The cells were then washed two times with fresh DMEM complete medium and replated into nocodazole free fresh medium and pulsed for 15 minutes with 10 µM EdU (Thermo Scientific) at 8 hr, 10 hr and 12 hr time points and fixed in 3.7% formaldehyde, and washed in PBS prior to EdU labeling by click chemistry. Cell populations were imaged with the IncuCyte ZOOM and the fraction of EdU-positive transfected (GFP-positive) cells was determined using the coincident analysis application within the IncuCyte software. For detection of DNA damage U2OS cells were seeded on glass coverslips and transfected with different Cdh1 constructs and histone H2B-GFP as a tracer. After 48 hours cells were fixed and permeablized with 0.5% Triton X-100 in PBS, washed and then blocked for 30 minutes at room temperature with 5% BSA in PBS. Cells were incubated with antibodies (1:500) in 5% BSA in PBST for 1 hour at room temperature. After washing the cells were incubated with Alexafluor secondary antibodies (1:500) in 5% BSA in PBST for 30 minutes at room temperature. DNA was counterstained with 1 µg/mL Hoechst 33342 and mounted with Fluorimount G (Southern Biotech). Cells were imaged using a Leica DM5500B fluorescent microscope as described previously. Images were analyzed and foci quantified with Cell Profiler software [85].

### Plasmids and Recombinant Proteins

HA-Cdh1 and different mutants of HA-Cdh1, cloned into pcDNA 3.1 were obtained from GenScript. The Cdh1 cDNA was amplified with PCR and the PCR products were subcloned into pCS2+ and pGEX-4T-1 vectors. Myc-Chk1, Myc-βTRCP1, Flag-βTRCP1, and HA-Plk1 were generated as described previously [84]. cDNAs for Chk1 and Chk1^L449R^ were a gift from Youwei Zhang and were subcloned into modified pCS2 vectors using Gateway cloning. Recombinant and *in vitro* translated proteins were produced as described [84].

### Antibodies

The following commercial antibodies, and the indicated concentrations, were used in this study. C-Myc (#E0115; 1:1000), Chk1 (G-4) (#H2714; 1:1000) and GST (Z-5) (#K0713; 1:1000) were purchased from Santa Cruz Biotechnology. M2 anti Flag Mouse antibody (#SLBT7654; 1:5000), cdc27 (AF3.1) (1:1000) and Actin (#087M4850; 1:10,000) were purchased from Sigma. Cdh1 (#CC43-100UG; 1:500) was purchased from Calbiochem. Cyclin A2 (BF683) (#6; 1:1000) and Phospho-Chk1Ser345 (133D3) (#15; 1:1000) were obtained from Cell Signaling. HA (#SJ254200; 1:1000) antibody was purchased from Biolegend. Plk1 (3F8) (#06050819; 1:500) was obtained from Enzo Life Sciences. Secondary antibodies for western blotting were purchased from LI-COR Biosciences. Anti-phospho-Histone H2AX (clone JBW301) (#2977883, 1:500) was purchased from EMD Millipore Corp. Alexa546-conjugated antibodies (#A11030) for immunofluorescence were purchased from Invitrogen.

### Western Blotting and Immunoprecipitation

Either HA-tagged Cdh1 and Myc-tagged Chk1 mutant or HA-tagged Cdh1 (or mutants) and Flag-βTRCP1 were expressed in 293T cells for 30 hours. Cells were treated with MG-132 (10 µM for 5 hr) prior to lysis. Cell extracts were generated in EBC buffer, 50mM Tris (pH 8.0), 120mM NaCl, 1% NP40, 1mM DTT, and protease and phosphatase inhibitors tablets (Thermo Fisher Scientific).For immunoprecipitation, equal amounts of cell lysates were incubated with the indicated antibodies conjugated to protein G beads (Invitrogen) or anti-HA beads (15µl per IP, Thermo Scientific) respectively from 4h to overnight at 4°C. The beads were then washed with EBC buffer including inhibitors. Binding to immobilized GST proteins was performed as described previously [40]. Immunoprecipitation samples or equal amount of whole cell lysates were resolved by SDS-PAGE, transferred to PVDF membranes (Milipore) probed with the indicated antibodies, and visualized with the LiCor Odyssey infra-red imaging system.

### In Vitro Kinase Assay

5 microgram indicated GST-Cdh1 fusion proteins was incubated with kinase reaction buffer (50 mM Tris pH 7.4, 10 mM MgCl_2_, 1 mM DTT, phosphatase inhibitors and 200 µM ATP) and 100 ng of Chk1 (Sigma) at 30°C for 45 minutes. Phosphorylated samples were immunoprecipitated on the glutathione beads (Life Technologies) and resolved by SDS-PAGE. Phosphorylation of GST-Cdh1 was detected by pIMAGO western-blot kit (Tymora Chemicals).

### Mass Spectrometry

HeLa cells were synchronized and harvested in G1/S boundary, after a 2 mM hydroxyurea (HU) treatment for 16 hr. Extracts were then prepared by resuspension in extract buffer (20 mM Tris-HCl, pH 7.2, 2 mM DTT, 0.25 mM EDTA, 5 mM KCl, 5 mM MgCl2) followed by two rounds of freeze-thaw and passage through a needle. Extracts were supplemented with ATP and an energy regenerating system. GST-Cdh1 was then incubated in extract for 1 hr at 30°C and then captured on Glutathione beads. After washing, the proteins were resolved on SDS-PAGE and visualized with Gelcode Blue (Pierce). Protein bands were digested with trypsin, extracted in 50% acetonitrile; 5% formic acid. After evaporation, peptides were resuspended in 1% acetic acid and analyzed on a Thermo Scientific Ultimate 3000 UHPLC + Orbitrap Elite hybrid Mass spectrometer. Phosphorylated GST-Cdh1 from *in vitro* kinase assay samples were also analyzed through Mass Spectrometry.

### In Vivo Ubiquitination

The *in vivo* ubiquitination assays were performed as described previously [40, 84]. Briefly, 293T cells were transfected with the constructs encoding HA-Cdh1 or HA-Cdh1 mutants, His-ubiquitin, Flag-βTRCP1 and Myc-L449R-Chk1 respectively. After a treatment with 20 µM MG132 for 5 hr, the cells were lysed with denaturing buffer (6 M guanidiniu-HCl, 0.1 M Na2HPO4/ NaH2PO4, 10 mM Tris-HCl pH 8.0, 10 mM Beta-Mercaptoethanol and 5 mM imidazole pH8.0), followed by sonication. After centrifugation, the lysates were collected and incubate with Ni-NTA agarose beads (QIAGEN) for 4 hr. His-ubiquitinated proteins were washed three times with denaturing buffer (8 M urea, 0.1 M Na2HPO4/ NaH2PO4, 10 mM Tris-HCl pH 6.3, 10 mM Beta-Mercaptoethanol and 0.2% or 0.1% triton-X-100) and eluted with elution buffer (150 mM Tris-HCl pH 6.7, 200 mM imidazole), resolved by SDS–PAGE and immunoblotted with the indicated antibodies.

### Clonogenic Survival Assay

HeLa and HCT116 cells were transfected with HA-Cdh1 (wild type or mutants) and pLKO.1 plasmid. 20 hr post transfection, cells were split into 6 well plates. Cells were then treated with 2 mM Hydroxyurea for 16 hr and released into fresh DMEM with puromycin (1µg/ml) followed by two washes with PBS. The cells were selected for 72 hr. After 8 days, the colonies were stained on the plates with crystal violet and counted. The images of the plates were taken in Li-COR and colonies were counted manually.

### IncuCyte Proliferation Assay for Live-Cell Analysis

Hela cells were transfected with the indicated DNAs and histone H2B-GFP as a tracer. 48 hours later the cells were placed in the IncuCyte Zoom System (Essen BioScience). Images were collected at 10x every 4 hours for 3 days. Proliferation of transfected cells was analyzed by quantifying GFP-positive nuclei in the Proliferation-Cell Count application of the IncuCyte software. Nuclear size was also determined using this application. Large nuclei were identified as having a size greater than or equal the largest nuclei frequently observed in control populations.

### Statistical Analysis

Statistical analyses were perforemed with GraphPad Prism Software using a 1- or 2-way ANOVA with Holm-Sidak’s or Dunnet’s post-test, respectively where appropriate (GraphPad Software, Inc.).

## Acknowledgements

This work was supported by The Ohio State University Comprehensive Cancer Center/Department of Radiation Oncology start-up funds and NIH grants R01 GM112895 and R01 GM108743 to MKS. The Orbitrap Elite instrument of the Lerner Research Institute Proteomics Core at the Cleveland Clinic Foundation was purchased via an National Institutes of Health shared instrument grant, 1S10RR031537-01. Research reported in the publication was supported by The Ohio State University Comprehensive Cancer Center and the National Institutes of Health under grant number P30 CA016058. The authors thank The Ohio State University Comprehensive Cancer Center’s Genomics Shared Resource for technical support. We also thank members of the Summers laboratory for insightful discussion and constructive comments on the manuscript. The content of this work is solely the responsibility of the authors and does not necessarily represent the official views of the National Institutes of Health.

## Author contributions

Conception and design: DP and MKS; Development of methodology: DP, AET, KD, AD, and MKS; Acquisition of data: DP, AM, AET, KD, AD BW; Analysis and interpretation of data, DP, AET, KD, BW, MV, and MKS; Writing, review, and/or revision of the manuscript: DP, MV and MKS. Study supervision: MKS. All authors read and approved the final manuscript.

**Figure EV1: Inhibition of Chk1 Enhances Stability of Endogenous Cdh1.**

HeLa cells were treated Chk1 inhibitor (where indicated) for 4 hr before adding 50 µg/ml cycloheximide (CHX) and HU. At the indicated time points, whole-cell lysates were prepared for immunoblot analysis.

**Figure EV2: Phosphorylation of Cdh1 by Chk1 Creates a Phosphodegron Recognized by SCF^βTRCP1^, Related to Figure 3.**

**(A)** 293T cells were transfected with the indicated HA-Cdh1 constructs together with Flag-βTRCP1. Cells were treated with proteasome inhibitor MG132 (10 µM) for 5 hr before immunoprecipitation to detect the interaction between HA-Cdh1 and Flag-βTRCP1proteins. A backround level of non-specifc interaction between Cdh1 proteins and the beads is observed in all conditions. Due to the lower amount of wild type Cdh1 the background in this band is only observed upon higher intensity scan (not shown).

**(B)** Constitutively active Chk1 promotes the interaction between Cdh1 and βTRCP1. Immunoblot analysis of immunoprecipitates and whole-cell lysates derived from 293T cells transfected with Flag-βTRCP1, HA-Cdh1 and Myc-Chk1^L449R^ (where indicated). 30 hr post-transfection, cells were treated with proteasome inhibitor MG132 (10 µM) for 5 hr and then harvested to do immunoprecipitation to detect the interaction between HA-Cdh1 and Flag-βTRCP1 proteins.

**(C)** 293T cells were transfected with the indicated HA-Cdh1 constructs together with Flag-βTRCP1 and Myc-Chk1^L449R^ (where indicated). Cells were treated with 50 µg/ml cycloheximide (CHX). At the indicated time points, whole-cell lysates were prepared for immunoblot analysis. The intensities of Cdh1 bands were normalized to actin, then normalized to the t=0 time point.

**Figure EV3: Chk1-Mediated Phosphorylation of Cdh1 Destabilizes the Cdh1-APC/C Interaction.**

*In-vitro* binding of wild-type or mutant Cdh1 proteins to the APC/C in G1/S extracts from HU arrested HeLa cells. Immunoblots represents both immunoprecipitated different HA-Cdh1 constructs with the relative inputs.

**Figure EV4: Model for the Role of Chk1-Mediated Cdh1 Phosphorylation on Replication Stress Response and S-Phase Entry.**

In early G1, APC/C^Cdh1^ maintains low levels of its substrates, including components required for Chk1 activitation. As cells near the G1/S transition APC/C^Cdh1^ activity diminishes (see text) and substrates accumulate, which allows activation of Chk1. Chk1 cooperates with Cyclin A and Plk1 to phosphorylate Cdh1 to create a phosphodegron which acts as a binding site for SCF^βTrCP^ leading to the ubiquitination and degradation of Cdh1. As cells leave S-phase, Plk1 targets Claspin for degradation to promote loss of Chk1 activity (see text). With decreasing Chk1 activity, Cdh1 stability increases and further inhibits additional activation of Chk1 by targeting Rad17 for degradation, thus promoting its own accumulation in G2.

**Figure EV5: USP37 Promotes Cdh1 Degradation via Chk1.**

HeLa cells were synchronized in mitosis with nocodazole following transfection with control (Ctrl) siRNA or USP37 siRNA during double thymidine. Cells were released from mitosis and whole-cell lysates were prepared at the times indicated to detect the impacts of USP37 knock down on both endogenous Cdh1 and pChk1 level in cells. The corresponding plot shows the Cdh1 band intensities at indicated time point normalized to actin and further normalized to the t=0 time point. Data are represented as mean +/- SD, n=2 biological replicates.

**Appendix Figure S1:** Mass spectrometry analysis of Chk1 mediated Cdh1 phosphorylation. Chymotryptic digests were performed on the Cdh1 protein in order to identify sites of phosphorylation. **(A)** The MS/MS spectra of a doubly charged SLSTKRSSPDDGNDVSPY + PO3 peptide was identified with a m/z ratio of 1002.9299 Da. The site of phosphorylation was identified to be either S131 or S133 by the mass of the y15 ion.

**(B)** The MS/MS spectra of a doubly charged SLSTKRSSPDDGNDVSPY + 2PO_3_ peptide was identified with a m/z ratio of 1042.9131 Da. The sites of phosphorylation was identified to be S131 and S133 by the mass of the b_3_ ion.

**(C)** The MS/MS spectra of a doubly charged KIpSKIPF peptide was identified with a m/z ratio of 456.7511 Da. The site of phosphorylation was identified to be S172 by the mass difference between the y_5_ and y_4_ ions

**Appendix Figure S2: (A)** Representative images of nuclear size, Histone H2B-GFP, related to Figure 5A and 5B. Scale bar = 50 µm

**(B)** Representative images of EdU positive cells (top panel), total transfected (green) cells (middle panel) and merged (bottom panel) are shown related to figure 5C. Scale bar = 50 µm

**(C)** Immunofluorescence analysis of DNA damage (γH2AX). Representative images of γH2AX focus formation in U2OS cells transfected with Histone H2B-GFP and the indicated Cdh1 constructs related to Figure 5D. Scale bar = 20 µm

**Appendix Figure S3: (A)** Mutation of the Chk1 phosphorylation sites in Cdh1 induces DNA damage. The number of 53BP1 foci in cells. Hela cells as in (Fig 5A) were analyzed for 53BP1 foci. The graph shows the number of 53BP1 foci per GFP-positive nucleus, n>500; *p< 0.05,, ****p<0.0001 were calculated with 1-way Anova with Dunnet’s post-test.

**(B)** Immunofluorescence analysis of DNA damage (53BP1). Representative images of 53BP1 focus formation in Hela cells transfected with Histone H2B-GFP and the indicated Cdh1 constructs. Scale bar = 20 µm

