## Supplementary figures and images for "Chk1 Phosphorylates Cdh1 to Promote SCF^βTRCP^-Dependent Degradation of Cdh1 During S-Phase"

### Supplemental Figures

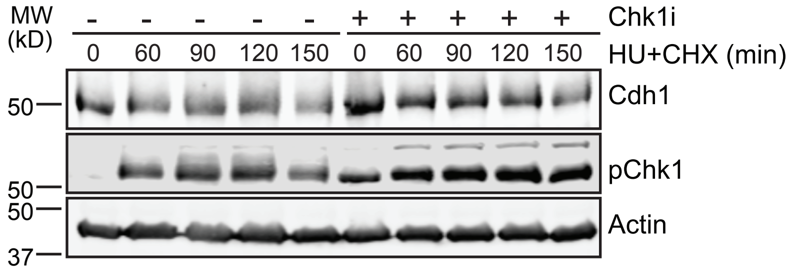

Pal et al., Figure EV2

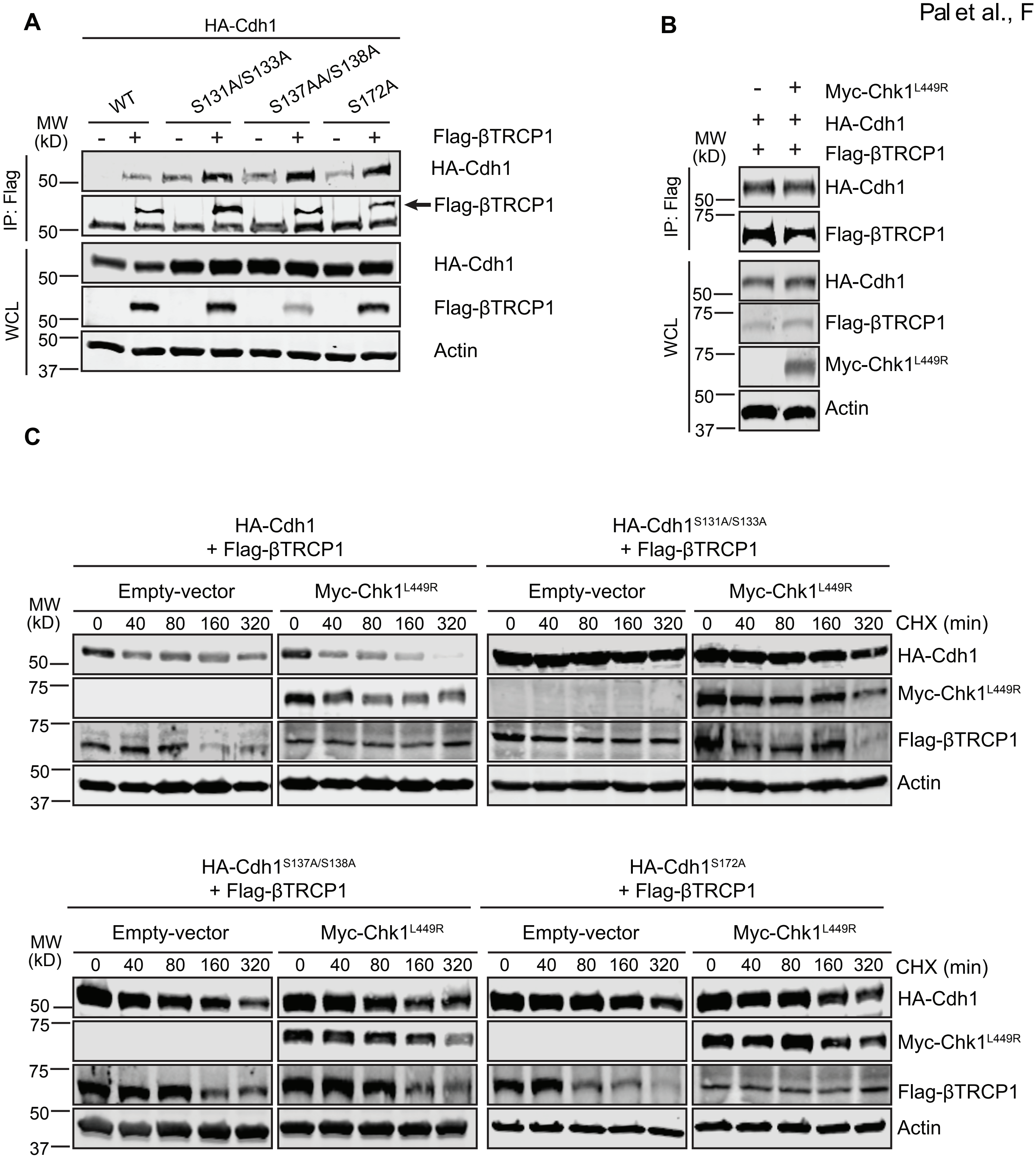

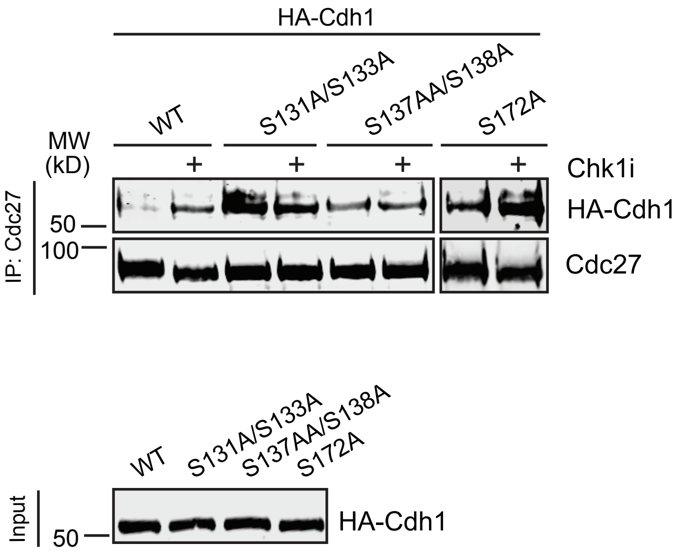

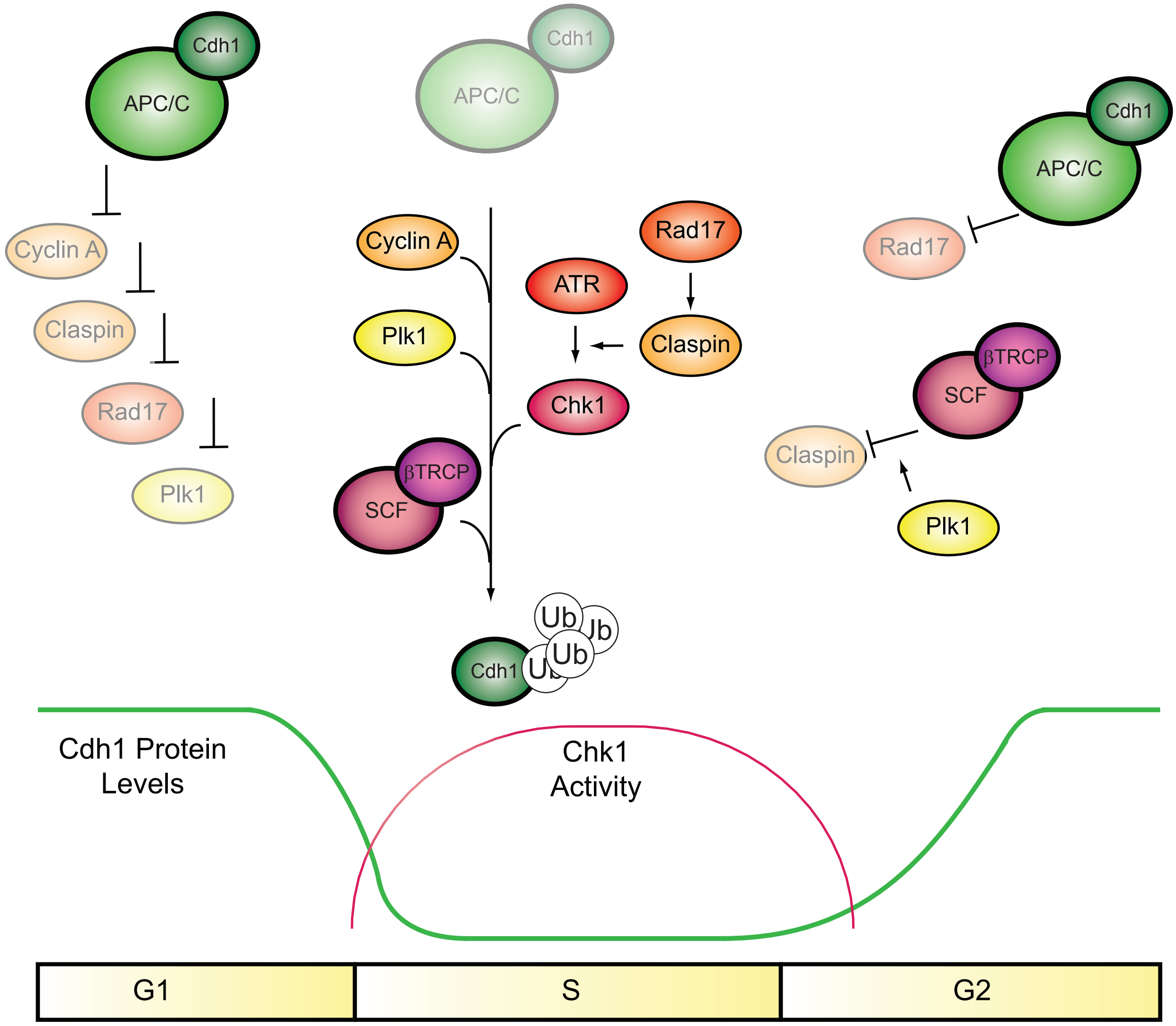

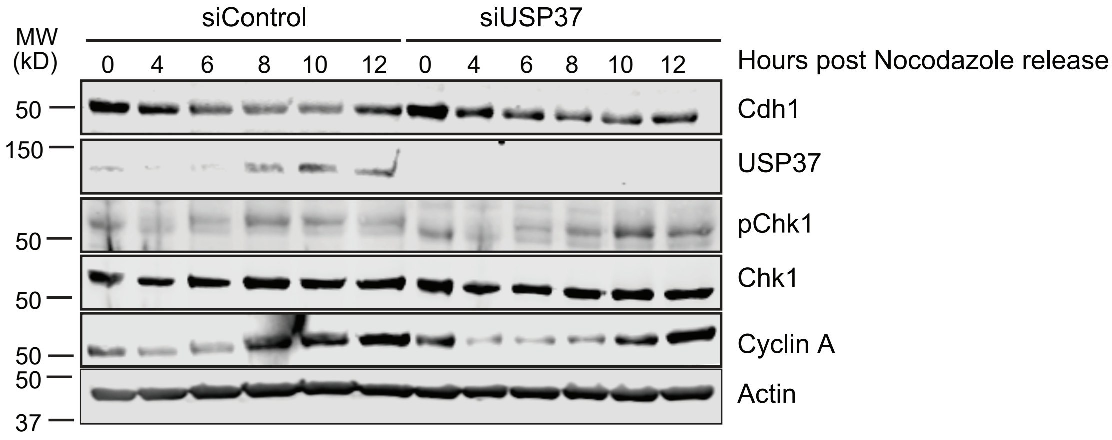

Pal et al., Appendix Figure S1

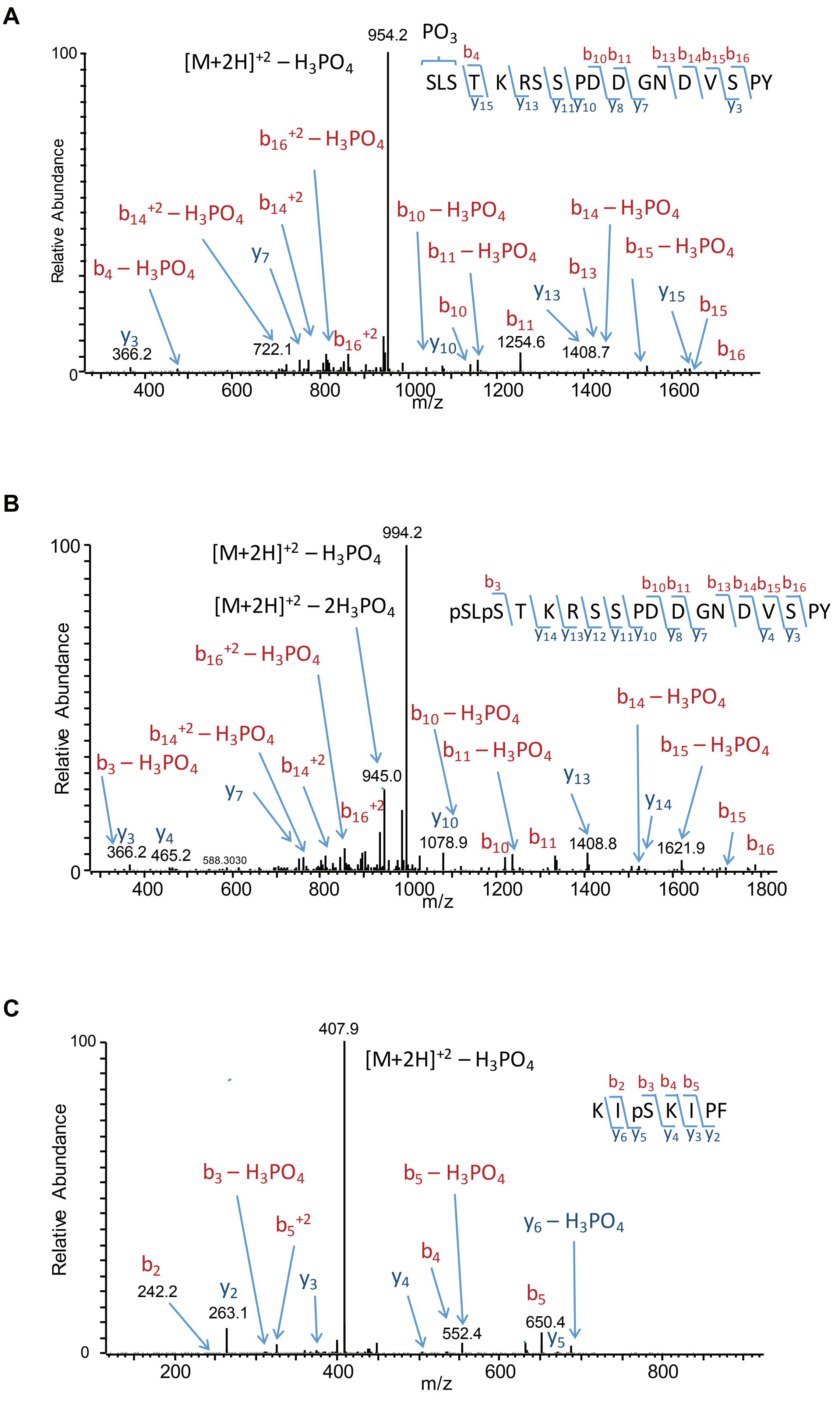

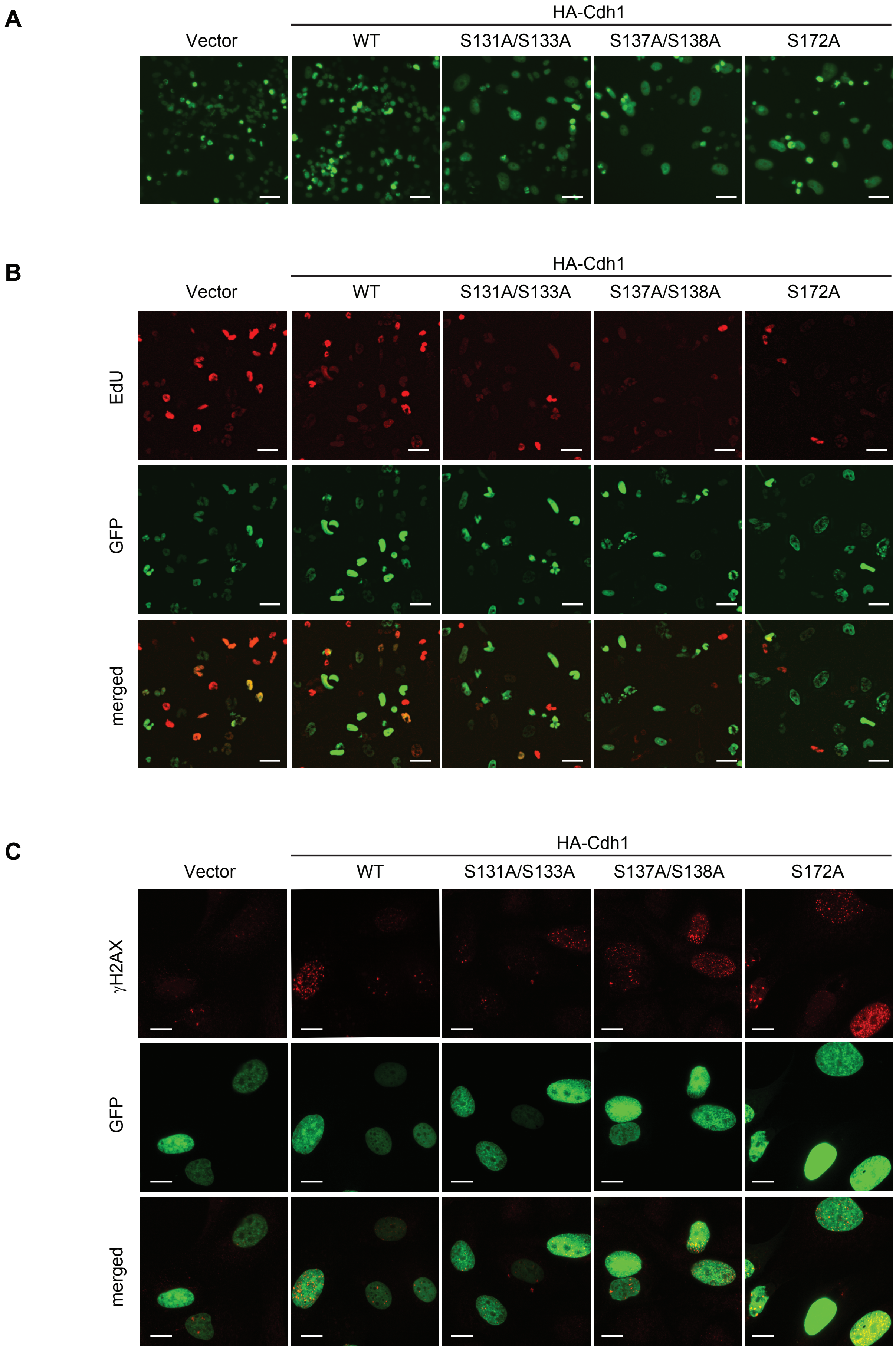

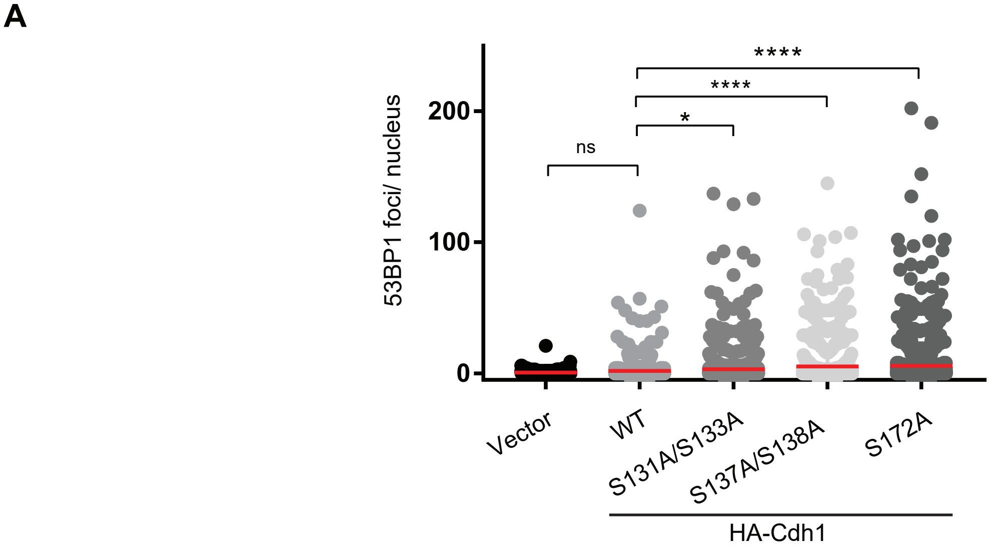

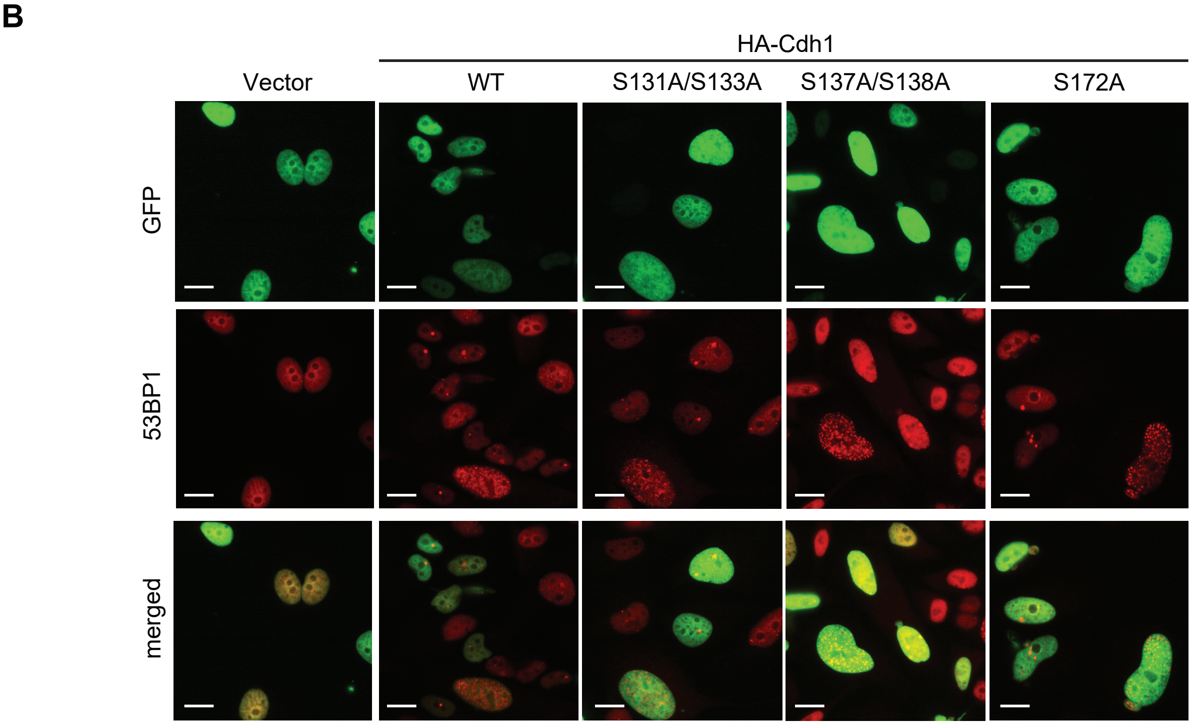
